# Q203 containing fully intermittent oral regimens exhibited high sterilizing activity against *Mycobacterium ulcerans* in mice

**DOI:** 10.1101/813253

**Authors:** Aurélie Chauffour, Jérôme Robert, Nicolas Veziris, Alexandra Aubry, Kevin Pethe, Vincent Jarlier

## Abstract

Buruli ulcer (BU), caused by *Mycobacterium ulcerans* is currently treated by a daily combination of rifampin and either injectable streptomycin or oral clarithromycin. An intermittent oral regimen would facilitate the treatment supervision. We first evaluated the bactericidal activity of newer antimicrobials against *M. ulcerans* using a BU animal model. The imidazopyridine amine Q203 exhibited high bactericidal activity whereas tedizolid (oxazolidinone close to linezolid), selamectine and ivermectine (avermectine compound) and the benzothiazinone PBTZ169 were not active. Consequently, Q203 was evaluated for its bactericidal and sterilizing activities in combined intermittent regimens. Q203 given twice a week in combination with one of the other long half-life compounds, rifapentine or bedaquiline, sterilized the mice footpads in 8 weeks, i.e. after a total of only 16 doses, and prevented relapse during a period of 20 weeks after stopping the treatment. These results are very promising for future intermittent oral regimens which would greatly simplify BU treatments in the field.

**Author summary:** The current treatment of Buruli ulcer (BU), infection caused by *Mycobacterium ulcerans* is based on a daily antibiotic combination of rifampin associated with streptomycin or clarithromycin. A shorter or intermittent treatment without an injectable drug would clearly simplify the management on the field. We evaluated the bactericidal activity of several new antimicrobials drugs in a mice model of BU and found that the Q203 exhibited the highest bactericidal effect. We subsequently identified new antibiotic combinations containing Q203 with high sterilizing activity when administrated twice a week for 8 weeks.

## Introduction

Buruli ulcer (BU), caused by *Mycobacterium ulcerans*, was only treated by surgery until 2004. The first medical treatment recommended by the World Health Organization (WHO). was a daily eight-week treatment based on an association of two antibiotics, rifampin (RIF), an oral ansamycin, and streptomycin (STR) an injectable aminoglycoside [1]. Currently a promising fully oral regimen combining RIF and clarithromycin (CLR), a macrolide compound [2,3] is tested clinically at a large scale (NCT01659437, clinicaltrials.gov).

The oral RIF-CLR combination has been promoted to suppress toxic effects and injections with aminoglycosides, resulting in better patients adherence and safety. Nevertheless, this combination is given daily during eight weeks. Shorter or intermittent treatment would facilitate adherence and the supervision by healthcare workers. For instance, many Buruli ulcer patients with small-to-moderate size wounds are on ambulatory care, and visit healthcare centres twice or three times per week for dressing changes, a rhythm that could allow receiving supervised intermittent antibiotic administration.

Our main objective was to identify alternative oral regimens active against BU by using a validated BU animal model. As a first step, we screened several new drug candidates for their *in vivo* bactericidal activity against *M. ulcerans*. Based on available data on activity against *M. tuberculosis* or *M. ulcerans*, the following compounds, with either short or long half-life, were selected as potentially interesting: selamectin (SEL) and ivermectin (IVE), two drugs from the avermectin family with antiparasitic properties [4–6]; tedizolide (TDZ) [7], a new oxazolidinone sharing the same mechanism of action as linezolid (LZD), a drug active against *M. ulcerans* [8] but exhibiting higher solubility and bioavailability than LNZ [9]; the 2-piperazino-benzothiazinone 169 (PBTZ), shown to be highly active against *M. tuberculosis* [10,11]; the imidazopyridine amine Q203 that targets the cytochrome bc1-aa3 respiratory terminal oxidase in *M. tuberculosis* and recently shown to prevent mortality and reduce CFU counts in the footpad of mice infected with *M. ulcerans* [12].

Because the first step screening demonstrated that Q203 was the compound with the highest bactericidal activity, and because of the long half-life of this compound, we measured in a second experiment the bactericidal and sterilizing activity of intermittent regimens containing Q203 combined with rifapentine or bedaquiline, other antibiotics with long half-life and already known to be active against *M. ulcerans*.

## Methods

### Infection of mice with *M. ulcerans*

Respectively 190 and 390 4 weeks-old female balb/c/j mice were used in the 1^st^ and the 2^nd^ experiment (Janvier Labs, Le Genest Saint-Isle, France). Mice were inoculated according to the Shepard method [13] in the left hind footpad with 0.03 ml of a bacterial suspension containing around 5 log_10_ Colony Forming Unit (CFU) of the *M. ulcerans* strain Cu001 (5.02 and 4.6 log_10_ in the 1^st^ and 2^nd^ experiment, respectively). This strain, isolated in 1996 from a Buruli ulcer patient in Adzopé, Ivory Coast [14], was kindly provided by the local laboratory without any identification data regarding the patient. The strain is susceptible to all drugs used in BU treatment and was maintained in our lab by regular passage into mice footpad.

### Treatment of mice

The treatment was initiated when the infection was well established, *i.e.* when the mice footpad swelling reached grades 2 (inflammatory swelling limited to the inoculated footpad) to 3 (inflammatory swelling involving the entire inoculated footpad) on a 4-grade ladder [15]. This stage of infection was reached six weeks after the inoculation. The mice were randomly allocated into eight groups (1^st^ experiment) and ten groups (2^nd^ experiment) using a randomization table (randomization.com).

The groups were as follows (drug, dosage, number of doses/week):

1^st^ experiment: one untreated control group of 30 mice and seven treated groups of 20 mice each treated with either tedizolide (TDZ) 10mg/kg 5/7, linezolide (LZD) 100mg/kg 5/7, selamectine (SEL) 12mg/kg 1/7, ivermectine (IVE) 1mg/kg 5/7, Q203 5mg/kg 5/7, 2-piperazino-benzothiazinone 169 (PBTZ) 25mg/kg 5/7 or, as controls, rifampin (RIF) 10 mg/kg 5/7, alone or combined with streptomycin (STR) 150 mg/kg.

2^nd^ experiment: one untreated control group of 27 mice, five groups of 27 mice each treated with monotherapy by RIF 10 mg/kg 5/7, rifapentine (RPT) 20mg/kg 2/7, bedaquiline (BDQ) 25 mg/kg 5/7 or Q203 5mg/kg 5/7 or 2/7; and 4 groups of 57 mice each treated with combined therapies Q203-RIF 5/7, Q203-RPT 2/7, Q203-BDQ 2/7 and RIF-clarithromycin (CLR) 100 mg/kg 5/7 as control.

LZD was purchased from Pfizer, France; SEL and IVE from Merck, France; RIF from Sandoz, France; STR from Panpharma, France; and CLR from Abbott, France. TDZ was kindly provided by MSD-MERCK group, BDQ by Janssen Pharmaceutica and PBTZ by Stewart Cole (Ecole Polytechnique Fédérale de Lausanne, Switzerland). Q203 was custom-synthetized at GVKBio.

Antibiotics were re-suspended in 0.05% agar-distilled water except for STR, which was diluted in normal saline, Q203 in 1% DMSO-20% TPGS (D-α-Tocopherol polyethylene glycol 1000 succinate) and PBTZ in a solution of 1% carboxyl-methylcellulose-1% Tween 80. BDQ was directly provided by Janssen Pharmaceutica in a 20% hydropropyl-β-cyclodextrin formulation.

Treatment regimens were administrated during 4 or 8 weeks in both experiments and all drugs were orally administered by gavage in a final volume of 0.2 ml, except STR, which was injected subcutaneously under the same volume.

### Assessment of *M. ulcerans* infection and effectiveness of treatment

Two methods were used for assessing the development of *M. ulcerans* infection and the effect of treatments: (i) clinical method by weekly evaluation of the lesion index as previously described [15], and (ii) bacteriological method by cultivating *M. ulcerans* from mice footpad. For CFU enumeration in footpads, mice were sacrificed by cervical dislocation as recommended by the European directive 2010/63 and the French decree n°2013-118. Mice footpads were removed aseptically and grinded in a Hank’s balanced salt solution under a final volume of 2 ml in an organ grinder (Octo Dissociator GentleMACS, Miltenyi®). Suspensions were then plated onto Lowenstein-Jensen (LJ) tubes containing Vancomycin 10µg/ml, Colistin 40µg/ml and B Amphotericin 10µg/ml to limit contamination. For untreated control groups, suspensions were serially diluted in 10-fold steps from pure to 10^−4^ and 0.1 ml of the dilutions were plated in duplicate on LJ-media whereas for the treated groups, the entire volume of the footpad suspension was plated onto 10 LJ-media with 0.2 ml each. All tubes were incubated at 30°C for 90 days.

In the 1^st^ experiment, lesion index of the footpads were measured during the 8 weeks period of treatment and CFU were numerated at week 4 and 8. In the 2^nd^ experiment, lesion index of the footpads were measured during the 8 weeks period of treatment and CFU were numerated at week 2, 4 and 8. Moreover, in the latter experiment, 30 mice that had been treated during 8 weeks with combined therapies were held without treatment during an additional period of 20 weeks to monitor relapses of *M. ulcerans* infection; lesion index were measured during this period and CFU were numerated at the end of the observation period.

### MIC determination

In order to assess a possible acquisition of resistance to the antibiotics used during treatment, MICs of the antibiotics used for the treatment were determined against the bacilli isolated from relapsing mice during the observation period and the initial strain Cu001. The strains were suspended in distilled water and the turbidity was adjusted to Mac Farland 3 (1 mg/ml). RIF and CLR were tested on a 7H11 + 10% OADC (Oleic-Acid-Dextrose-Catalase) medium (pH 7.4) and CLR was tested also on Mueller Hinton medium (pH 6.6). RIF was dissolved in dimethylformamide and CLR in distilled water, then twofold diluted in their own solvent and incorporated to the culture-media to obtain a final concentration ranging from 4 to 0.12 μg/ml. 0.1 ml of two distinct bacilli suspensions (pure and 10^−2^) were plated onto drug-containing-media and drug-free media used as a growth control. All media were incubated at 30°C and examined after 60 and 90 days.

### Statistical analysis

The Mann-Whitney test was used to analyze the results. A p-value<0.05 was considered as statistically significant. A regimen was considered to be bactericidal if its mean value of CFU per footpad was significantly lower than the mean value of CFU per footpad of the untreated group.

### Ethic statement

Experiment project was favorably evaluated by the ethic committee n°005 Charles Darwin localized at the Pitié-Salpêtrière Hospital and clearance was given by the French Ministry of Education and Research under the number APAFIS#9576-20170301171176185 v2. Our animal facility received in April 27^th^ 2017 authorization to carry out animal experiments with the license number C-75-13-01. The persons who carried out the animal experiments had followed a specific training recognized by the French Ministry of Education and Research.

## Results

### 1^st^ experiment

#### Evolution of the footpad lesions (Fig 1)

The mean lesion index (MLI) was 2.8 at the start of the treatment. Footpads of untreated control mice swollen from MLI 2.8 to 4 after 2 weeks and mice had to be sacrificed at week 4 due to advanced lesions. MLIs in RIF containing control groups increased to 3.8 after one week and then decreased to remain stable at 3.4 and decreased to 2 in RIF-STR group. MLIs in the SEL, IVE, TDZ and PBTZ groups continued to increase and mice had to be sacrificed at week 4 due to advanced lesions. In contrast, MLI in the Q203 treated group decreased to 1.9 after 1 week of treatment and to 1.2 after 8 weeks.

**Fig 1.**
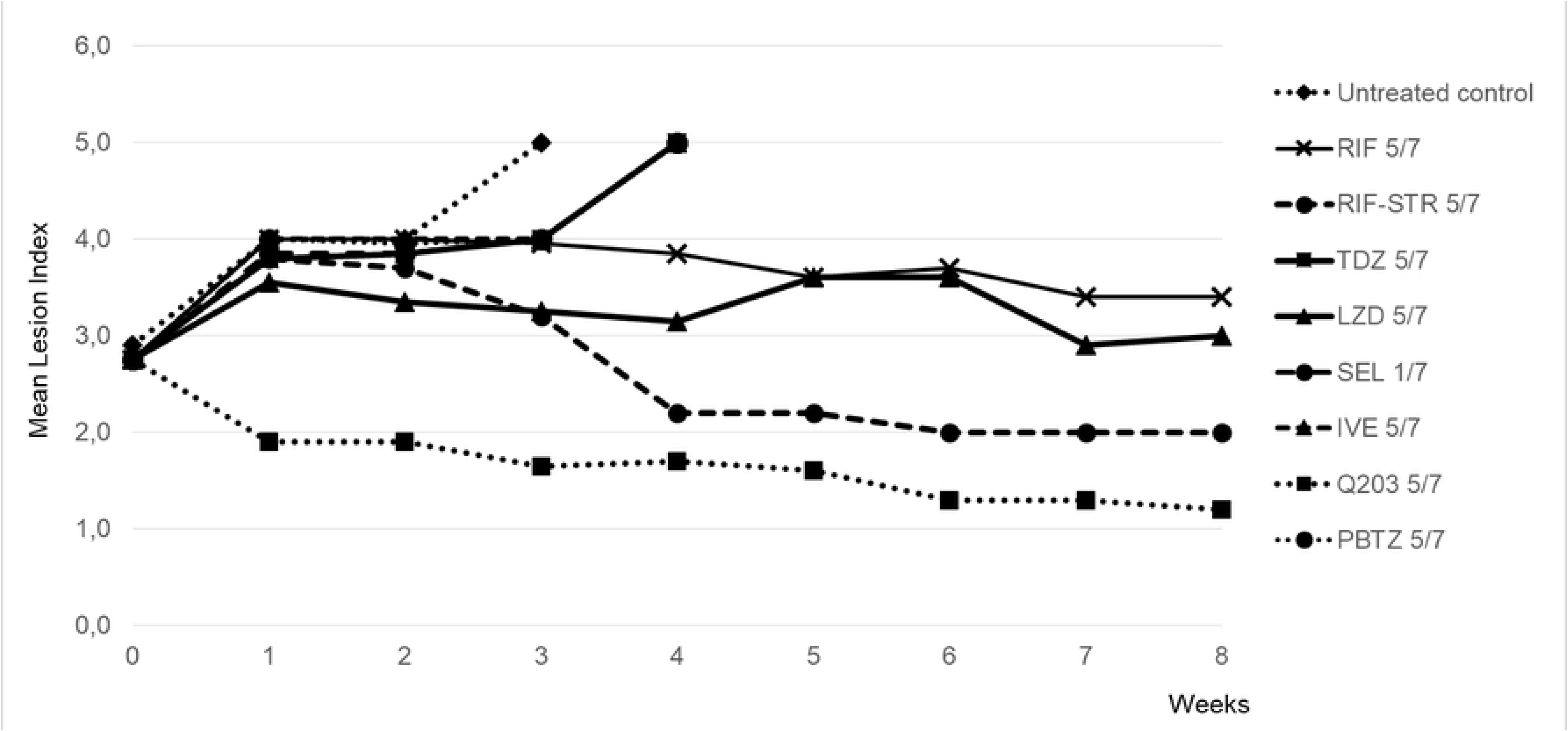
First experiment: Evolution of the mean lesion index of the swelling footpad of mice infected with *M. ulcerans* during 8 weeks of treatment. Doses were as follow: Rifampin (RIF) 10mg/kg; Streptomycine (STR) 150mg/kg; Tedizolide (TDZ) 10mg/kg; Linezolide (LZD) 100mg/kg; Selamectine (SEL) 12 mg/kg; Ivermectine (IVE) 1 mg/kg; Q203 5mg/kg; PBT169 25mg/kg

#### Evolution of the CFU counts (Table 1)

All untreated mice had culture-positive footpads at the time of treatment start with a mean of 6.93 ± 0.20 log_10_ CFUs, value that remained unchanged at week 4. All mice remained culture positive after 4 weeks of treatment in the RIF control-groups but the mean CFU counts was significantly lower (p<0.05) than that in the untreated mice group (4.58 ± 1.29 log_10_ for RIF and 2.15 ± 1.20 log_10_ for RIF-STR, respectively). After 8 weeks of treatment, only 2 out of 8 mice in the RIF groups remained culture positive with a low mean CFU counts (<1 log_10_). After 4 weeks of treatment, all the mice were culture positive in the TDZ, SEL, IVE and PBTZ groups, with CFU counts not statistically different from those in the untreated control group. Due to advanced footpads lesions and mortality of mice, the evaluation point initially planned at week 8 was cancelled for these groups. After 4 weeks of treatment, 3/10 mice were already culture negative in the Q203 treated group with a low mean CFU count (1.14 ± 1.30 log_10_), i.e. different (p< 0.001) from those in the RIF-containing group. After 8 weeks of treatment, all mice were culture negative in the Q203 group.

**Table 1.** *First experiment*: Results of footpad cultures during the treatment of mice infected with *M. ulcerans*.

| Regimen<sup>a</sup> | Results during treatment |  |  |  |  |  |
| --- | --- | --- | --- | --- | --- | --- |
|  | D0 |  | 4w |  | 8w |  |
|  | Culture positivity rate | Mean (±SD) CFU per group | Culture positivity rate | Mean (±SD) CFU per group | Culture positivity rate | Mean (±SD) CFU per group |
| Untreated control | 10/10 | 6,93±0,20 | 13/13<sup>b</sup> | 6,81±0,36 |  |  |
| RIF 5/7 |  |  | 9/9<sup>d</sup> | 4,58±1,29 | 2/8<sup>d</sup> | 0,47±1,16 |
| RIF-STR 5/7 |  |  | 8/8<sup>e</sup> | 2,15±1,20 | 3/9<sup>e</sup> | 0,57±0,90 |
| TDZ 5/7 |  |  | 7/7<sup>f</sup> | 6,35±0,59 | <sup>g</sup> |  |
| LZD 5/7 |  |  | 10/10 | 2,83±0,62 | 9/10 | 1,87±1,11 |
| SEL 1/7 |  |  | 4/4<sup>h</sup> | 6,24±0,82 | <sup>g</sup> |  |
| IVE 5/7 |  |  | 8/8<sup>i</sup> | 6,30±0,82 | <sup>g</sup> |  |
| Q203 5/7 |  |  | 7/10 | 1,14±1,30 | 0/10 |  |
| PBTZ169 5/7 |  |  | 7/7<sup>j</sup> | 7,08±0,64 | <sup>g</sup> |  |

<sup>a</sup>: treatment began 6 weeks after inoculation of 5.02 log<sub>10</sub> per footpad when the infected swelling footpads reached a lesion index between 2 and 3.
Drugs were administered 5 times a week except for the selamectine group with 1 time a week. Dosages were as follow: Rifampin (RIF) 10mg/kg; Streptomycine (STR) 150mg/kg; Tedizolide (TDZ) 10mg/kg; Linezolid (LZD) 100mg/kg; Selamectine (SEL) 12 mg/kg; Ivermectine (IVE) 1 mg/kg; Q203 5mg/kg; PBT169 25mg/kg.
<sup>b</sup>: due to advanced lesion, all mice from the untreated control group were in fact sacrificed at 3 weeks; despite that, footpad cultures were contaminated for 7 out 20 mice.
<sup>d</sup>: 3 mice died in the RIF groups due to an accident of gavage.
<sup>e</sup>: 3 mice died in the RIF-STR groups due to an accident of gavage.
<sup>f</sup>: footpad cultures were contaminated due to advanced lesion for 3 mice.
<sup>g</sup>: cultures at week 8 were not performed due to advanced necrotized lesion in footpads.
<sup>h</sup>: 5 mice died from *M.ulcerans* infection and footpad cultures were contaminated due to advanced lesion for 1 mouse.
<sup>i</sup>: 1 mouse died from *M.ulcerans* infection and footpad cultures were contaminated due to advanced lesion for 1 mouse.
<sup>j</sup> : footpad cultures were contaminated due to advanced lesion for 3 mice.

### 2nd experiment

#### Evolution of the footpad lesions (Fig 2)

The MLI was 3 at the start of the treatment. Footpads of untreated control mice swollen from MLI 3 to 4 after 2 weeks and mice had to be sacrificed at week 4 due to advanced lesions. MLIs in RIF, RPT and BDQ treated groups increased to 3.8-4 after one week of treatment and then decreased to reach at week 8 the values 3.6 (BDQ), 3 (RIF) and 2.7 (RPT). In the Q203 treated groups (5/7 or 2/7), MLIs slightly increased to 3.5 after one week of treatment and then rapidly decreased to reach 1.6-1.7 at week 8. MLI in the group treated with RIF-CLR increased to 4 after one week of treatment and decreased smoothly to reach 1.4 at week 12, remained stable until week 20, but increased again afterwards to reach 2.6 at week 28. MLI in the group treated with Q203-BDQ slightly increased to 3.6 after one week of treatment and decreased smoothly thereafter to reach 1.4 at week 12, a level stable till week 28. MLIs in the groups treated by Q203 combined with RIF or RPT rapidly decreased to 1.2-1.3 at week 8, and remained at this level till week 28.

**Fig 2.**
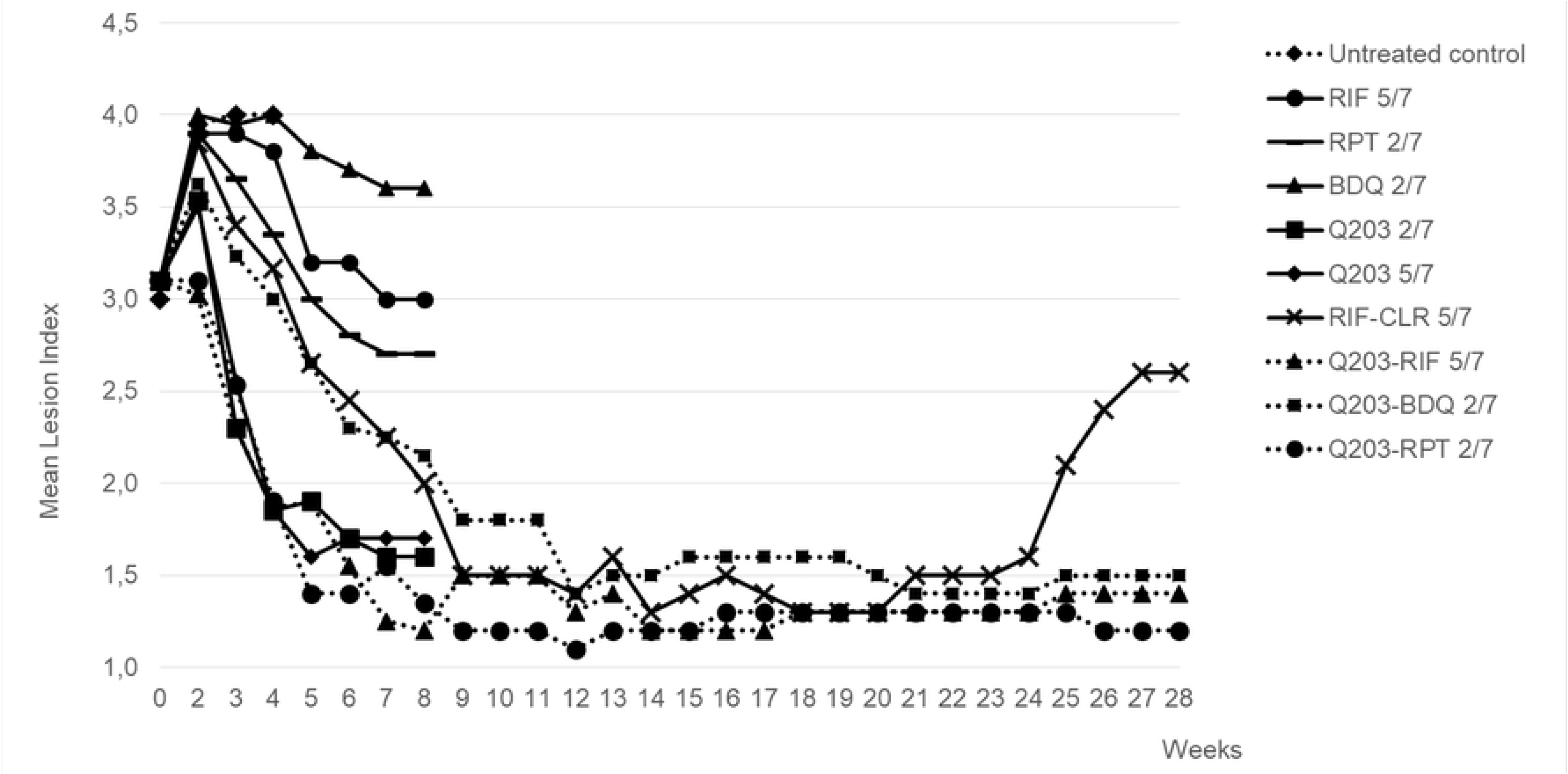
Second experiment: Evolution of mean lesion index of the swelling footpad of mice infected with *M. ulcerans* during 8 weeks of treatment and during 20 weeks of relapse observational period. Dosages were as follow: Rifampin (RIF) 10mg/kg; Rifapentine (RPT) 20mg/kg; Bedaquiline (BDQ) 25mg/kg; Q203 5mg/kg; Clarithromycin (CLR) 100mg/kg

#### Evolution of the CFU counts (Table 2)

All untreated mice had culture-positive footpads at the time of treatment start with a mean of 6.87 ± 0.10 log_10_ CFUs, value that remained unchanged at week 2 and 4. All treated mice remained culture positive after 2 weeks of treatment but CFU counts were significantly lower in all treated groups compared to those in untreated group. Moreover, CFU counts were lower (p≤0.01) in RPT, Q203 and in the 4 combined treatment groups than in RIF and BDQ groups. After 4 weeks of treatment, part of the mice became culture negative especially in the groups treated with Q203-RPT and Q203-BDQ were CFU counts were very low (~0.2 log_10_). CFU counts at 4 weeks remained lower (p<0.05) in RPT and Q203 than in RIF and BDQ groups and were lower in Q203-RPT and Q203-BDQ groups than in Q203-RIF and RIF-CLR groups. After 8 weeks of treatment, all of the mice treated with Q203 alone or combined with RIF 5/7 or RPT 2/7 or BDQ 2/7 became culture negative. Few mice remained culture positive in RIF, RPT and RIF-CLR groups with very low mean CFU counts (<0.5 log_10_) but all were still culture positive in the BDQ group (1.28 ± 0.93 log_10_).

**Table 2.** Results of footpad cultures during the treatment of mice infected with *M. ulcerans* (2, 4 and 8 weeks) and relapse rate after treatment completion.

| Regimen<sup>a</sup> | Results during treatment |  |  |  |  |  |  |  |  |  |
| --- | --- | --- | --- | --- | --- | --- | --- | --- | --- | --- |
|  | D0 |  | 2w |  | 4w |  | 8w |  | 28w |  |
|  | Culture positivity rate | Mean (±SD) CFU per group | Culture positivity rate | Mean (±SD) CFU per group | Culture positivity rate | Mean (±SD) CFU per group | Culture positivity rate | Mean (±SD) CFU per group | Culture positivity rate | Mean (±SD) CFU per group |
| Untreated control | 9/9 | 6.87±0.10 | 8/8<sup>b</sup> | 7.00±0.16 | 6/6<sup>b</sup> | 6.84±0.35 |  |  |  |  |
| RIF 5/7 |  |  | 9/9 | 5.71±0.65 | 6/6<sup>c</sup> | 2.78±0.81 | 2/8<sup>c</sup> | 0.44±0.81 |  |  |
| RPT 2/7 |  |  | 8/9 | 4.18±1.71 | 3/9 | 0.63±1.03 | 1/9 | 0.20±0.59 |  |  |
| BDQ 2/7 |  |  | 7/7<sup>d</sup> | 6.12±0.31 | 7/7<sup>d</sup> | 5.48±0.14 | 5/5<sup>d</sup> | 1.28±0.93 |  |  |
| Q203 2/7 |  |  | 9/9 | 4.88±0.0.42 | 5/9 | 1.42±1.21 | 0/9 |  |  |  |
| Q203 5/7 |  |  | 9/9 | 4.90±0.29 | 7/9 | 0.74±0.79 | 0/9 |  |  |  |
| RIF-CLR 5/7 |  |  | 9/9 | 4.42±0.55 | 6/7<sup>e</sup> | 1.15±0.84 | 1/8<sup>e</sup> | 0.22±0.63 | 8/26<sup>e</sup> | 0.87±1.36 |
| Q203-RIF 5/7 |  |  | 9/9 | 4.52±0.36 | 9/9 | 1.20±1.04 | 0/8<sup>f</sup> |  | 0/30 |  |
| Q203-BDQ 2/7 |  |  | 9/9 | 4.65±0.75 | 2/9 | 0.17±0.51 | 0/9 |  | 0/30 |  |
| **Q203-**<br>**RPT 2/7** |  |  | 9/9 | 4.40±0.78 | 2/9 | 0.25±0.49 | 0/9 |  | 0/30 |  |

<sup>a</sup>: treatments was began 6 weeks after inoculation of 4.6 log<sub>10</sub> per footpad when the infected swelling footpads reached a lesion index between 2 and 3. Drugs were administered 2 or 5 times a week and dosages were as follow: Rifampin (RIF) 10mg/kg; Rifapentine (RPT) 20mg/kg; Bedaquiline (BDQ) 25mg/kg; Q203 5mg/kg; Clarithromycin (CLR) 100mg/kg.
<sup>b</sup>: footpad cultures were contaminated due to advanced lesion for 1 mouse in the untreated control group 2 weeks and 1 mouse in the group 4 weeks.
<sup>c</sup>: footpad cultures were contaminated due to advanced lesion for 3 mice in the RIF group 4 weeks and 1 mouse in the group 8 weeks.
<sup>d</sup>: footpad cultures were contaminated due to advanced lesion for 1 mouse in the BDQ group 2 weeks, 2 mice in the group 4 weeks and 2 mice in the group 8 weeks; 2 mice died due to an accident of gavage in the group 8 weeks.
<sup>e</sup>: footpad cultures were contaminated due to advanced lesion for 2 mice in the RIF-CLR group 4 weeks and 3 mice in the relapse observation group; 1 mouse died due to an accident of gavage in the group 8 weeks and in the relapse observation group, respectively.
<sup>f</sup>: 1 mouse died due to an accident of gavage in the Q203-RIF group 8 weeks.

During the 20 weeks observation period after stopping the treatment, no bacteriological relapse was observed in the 3 groups treated with Q203 combinations whereas 8 / 26 mice treated with RIF-CLR relapsed with a mean CFU count of 0.87 ± 0.93 log_10_.

#### MIC of *M*. *ulcerans* bacilli recovered from relapsing mice

MICs remained unchanged against bacilli isolated from the 8 relapsing mice in the RIF-CLR treated group when compared to initial MICs against *M*. *ulcerans* Cu001 *i.e.* 0.5-1 μg/ml for RIF, and 0.5 μg/ml for CLR (for the latter, same value on 7H11 and MH media).

## Discussion

Although Buruli ulcer can be successfully treated by a two-month antibiotic combination regimen administered daily, new shorter and/or intermittent regimens, would greatly simplify treatment management in the field.

Our 1^st^ screening experimental in vivo study aimed at identifying newer bactericidal drugs. Indeed, available data on activity of several new drugs against *M. tuberculosis* or *M. ulcerans* justifying a systematic evaluation in a BU mouse model that has been successfully used for many years for this purpose [8]. Ivermectine, selamectine and tedizolid were not bactericidal after 4 weeks of treatment and failed to prevent mortality in 8 weeks. The doses used in our experiment were drawn from available pharmacokinetic data. Ivermectine, a long lasting drug in human (half-life 15-19h) and in mice (9h) was shown when used in mice at a dose of 0.2mg/kg to yield serum concentration lower than that obtained in human at standard therapeutic dose [12]. We therefore used a higher dose (1 mg/kg) that, still, failed to control infection in our model. Same unfavorable result was obtained with selamectine used at a dose of 12mg/kg as proposed in previous publication [19]. However, it has been suggested that these two avermectin compounds might be used safely at higher doses [12], which could be evaluated in future studies. The dose of tedizolid TDZ used in the present study, 10 mg/kg, was shown to yield in mice pharmacokinetics close to that obtained in human at the therapeutic dose of 200mg [20-22]. Contrasting with the deceiving results obtained with tedizolid, and as reported in a previous work [8], a marked bactericidal activity was obtained with linezolid, an oxazolidinone included in the experiment as a positive control for tedizolid. Surprisingly, PBTZ169 was not bactericidal in our BU model at 25 mg/kg, a dose shown to be active against *M. tuberculosis* in mice [15]. Yet *M. ulcerans*, as *M. tuberculosis*, carries a cysteine at the position 387 in DprE1, that codes for the target of benzothiazinones, but not a serine or an alanine, which has been shown to confer a natural resistance to PBTZ in *M. avium* or *M. aurum* [16]. Thus, the reason for the disappointing result obtained with PBTZ in our BU model is unclear.

RIF and RIF-STR were highly bactericidal as in all our preceding works [17]. Q203 drastically reduced the lesion index and CFU counts after 4 weeks of treatment and all mice became culture negative after 8 weeks. These results obtained with Q203 were significantly better than those obtained with the historical positive control RIF alone and even with the reference combination regimen RIF-STR.

Although drugs with low bactericidal activity when given in monotherapy might be of interest when used in combination with other drugs, and since our goal was to obtain the most effective combination regimen, we selected for the 2^nd^ experiment combinations of drugs shown to be highly active separately. The results of this experiment demonstrated that regimens combining Q203 with RIF or RPT or BDQ were not only bactericidal, making all the mice culture negative after 8 weeks of treatment, but also sterilized the mice footpads and prevented relapse during an observation period of 20 weeks after stopping the treatment. Importantly, these impressive results were obtained when administrating twice weekly during 8 weeks, i.e. after a total of only 16 doses, the combinations of Q203 with either RPT or BDQ, all long lasting drugs (serum half-life in mice after single dose: Q203 23 h, RPT 25 h, BDQ 53 h). Recently, regimens combining RPT), with CLR or BDQ, administrated twice weekly during 8 weeks were found as bactericidal and as sterilizing as daily RPT-CLR regimen [4].

The fact that, in the present work, few bacilli were still found by culture in 1/8 mice after 8 weeks of treatment with RIF-CLR 5/7, and that 8/26 mice relapsed within 20 weeks after the end of this regimen was surprising since bactericidal and sterilizing activity of such regimen was shown in two previous studies [17,18]. The susceptibility to RIF and to CLR of the bacilli isolated from relapsing mice was unchanged, ruling out the selection of resistant mutants during treatment. Relapses could be explained by an unusually high inoculum reached in the present work when compared to those in the previous studies, i.e. 3-4 times higher at the start of treatment and 4-10 times higher after 2-4 weeks in untreated mice. Nevertheless, this fact strengthens further the good results obtained with the Q203 combined regimens.

New drugs are rare for the treatment of BU. The last active new marketed drug was BDQ, initially experimented with success in tuberculosis, and found later on to be very active against *M.ulcerans* in BU animal model [17]. Therefore, the excellent *in vivo* activity of Q203 against *M. ulcerans* constitutes a step forward. The reason of this promising result has been elucidated in a recent study: reductive evolution in most strains of this species led to hyper susceptibility to Q203 by eliminating alternate terminal electron acceptors and thus making the target cyt-bc1:aa3 crucial for survival [12].

Triple combinations of Q203, administered at the higher dose of 10 mg/kg, either with RPT and clofazimine, RPT and BDQ or BDQ and clofazimine, as well as quadruple combination of these four drugs, were recently found to be sterilizing after 2 weeks of daily treatment in BU animal model [19], leading to the conclusion that targeting the *M. ulcerans* respiratory chain with several drugs is an efficient strategy for designing new shorter treatments of BU. In the present study, we demonstrated that double combinations of Q203 at 5 mg/kg with either RPT or BDQ, thanks to the long half-life and good bio availabilities of these drugs, provided very promising results for future fully oral intermittent regimens which would greatly simplify BU treatments in the field.

